# The Induction of Larval Resource Preference in Heterogeneous Habitats

**DOI:** 10.1101/274852

**Authors:** Vrinda Ravi Kumar, Swastika Issar, Deepa Agashe

## Abstract

Animals often have to evaluate and choose between multiple food sources in their habitat, and these potentially complex decisions can have a large impact on their fitness. Among other factors, previous experience with an alternative resource can significantly increase subsequent preference for the resource ("induction of preference"). Such induction of resource preference is particularly relevant in spatially or temporally heterogeneous habitats. Although most mobile species – especially generalists – probably frequently encounter habitat heterogeneity, the impact of preference induction on individual behaviour and fitness in heterogeneous habitats is poorly understood. We analysed larval preference induction in wheat-adapted generalist red flour beetles (*Tribolium castaneum*) under three types of habitat heterogeneity. We first analysed the induction of larval preference for novel resources (other cereal flours) under temporal heterogeneity, exposing larvae to new resources during development. We found that larvae preferred a new resource if they experienced it recently, but that the magnitude of induction varied across resources. Interestingly, we also observed specific induction for a homogenous mix of wheat and a novel resource, with larvae preferring the mix over either pure resource. To analyse induction under spatial heterogeneity, we placed beetle eggs in one of two alternative resource patches and tested the preference of emerged larvae. Unexpectedly, hatching into a novel resource did not always induce preference. Finally, we found that induction of preference for new resources could be maladaptive for larval development. Together, our work demonstrates that experience-based plasticity of larval resource choice may strongly impact larval preference and fitness in heterogeneous habitats.

## INTRODUCTION

Insects occupy a large diversity of dietary niches, and a substantial body of work has thus focused on understanding insect resource choice. Changes in resource preference can facilitate dietary niche shifts (Agosta 2006), potentially leading to divergence in host-associated traits (Singer & McBride 2010). Thus far, most studies of resource choice in holometabolous insects have analyzed only adult preference, since it is often well correlated with larval performance (Dethier 1959; Jaenike 1978; Gripenberg et al. 2010). However, many larvae also exhibit resource choice (Jermy et al. 1968; Berdegue & Trumble 1996; Bernays & Weiss 1996; Gamberale-Stille et al. 2014) and are capable of exploring different habitat patches over smaller spatial scales (Abbott & Dukas 2016). Thus, even though adult oviposition preference often determines larval resource use, it is important to study larval choice in spatially heterogeneous environments, where larval preference may also contribute substantially to larval fitness. Under such conditions, it is possible that resource choice is not the sole domain of the adult, and may instead be shared across stages (Wiklund 1975; Chew 1977; Berdegue et al. 1998; Gamberale-Stille et al. 2014; Abbott & Dukas 2016). However, because larval preference is relatively understudied, we do not know the impact of larval preference on larval fitness in heterogeneous environments.

What factors shape larval preference in heterogeneous habitats? A large body of work suggests that prior experience with alternative resources can strongly shape larval preference. Many Lepidopteran larvae show “induction of preference”: an increased preference for host plants that they have experienced previously (Jermy et al. 1968; Bernays & Weiss 1996; Carlsson et al. 1999; del Campo et al. 2001; Henniges-Janssen et al. 2014; Soler et al. 2012; Gretes et al. 2016). A particularly striking example is the induction of preference in *Manduca sexta*. While naïve larvae can feed and survive on many host plants, larvae that feed on solanaceous foliage become specialized and reject other hosts (Schoonhoven 1967; Yamamoto 1974; del Campo & Renwick 2000). Similar induction of preference also occurs in some non-Lepidopteran insects (Phillips 1977). Although preference induction thus seems to be common across insects, little is known about its occurrence in natural conditions and its impact on insect fitness. Previous studies suggest that the induction of preference could lead to increased acceptability of the inducing host, improving subsequent insect performance on it through higher feeding and/or physiological acclimation (Schoonhoven & Meerman 1978; Scriber 1979; Karowe 1989; Agrawal et al. 2002). However, these studies focus on mechanistic aspects of experience-mediated changes in food acceptability and digestive efficiency, leaving fundamental questions about the induction of preference under resource heterogeneity unanswered. For instance, we do not know the impact of experience with temporally or spatially heterogeneous resources on larval preference, and consequently on larval resource use. This is particularly important since most animals probably encounter significant habitat heterogeneity over their lifespan. Larval movement in a heterogeneous habitat could result in larvae simultaneously or sequentially experiencing different resources, but how does the induction of preference operate in such cases? For instance, does the duration of exposure to a resource impact preference, or is preference determined during a specific developmental period? Does the magnitude of induction vary depending on available resources? Does induction by the natal resource dictate larval resource choice in a patchy habitat (as is often assumed), or does the presence of alternative resources decrease the impact of the natal experience on larval choice? Finally, is the induction of larval preference adaptive?

To address these questions, we tested the occurrence and strength of induction of larval resource preference in a generalist insect pest of stored grain flour – the red flour beetle *Tribolium castaneum* (Sokoloff 1974; Ziegler 1976). Flour beetles are strong dispersers and frequently colonize grain warehouses (Naylor 1961; Hagstrum & Gilbert 1976; Ziegler 1976; Ziegler 1977). Resource availability within storage warehouses fluctuates over time, and beetles often move across flour patches (Ziegler 1976; Campbell & Hagstrum 2002), experiencing several resources in their lifetime and potentially exerting resource choice. Prior work suggests that *T. castaneum* larval preference can oppose adult resource preference: given a choice between the ancestral wheat resource and a suboptimal novel resource (corn), *T. castaneum* larvae favoured wheat, while adult females preferentially oviposited in corn (Agashe & Bolnick 2012). At high density, larvae raised in a heterogenous wheat-corn patchy habitat also preferred corn more strongly than larvae from a homogenous wheat habitat (Parent et al. 2014), suggesting that prior experience in a heterogeneous habitat alters larval resource choice in the face of competition. Thus, *T. castaneum* larval choice may determine larval resource use in heterogeneous environments. Here, we examined the role of larval choice and the impacts of resource heterogeneity on larval resource preference and fitness in *T. castaneum.* We first tested whether individual *T. castaneum* larvae show induction of preference for four novel resources, given temporal heterogeneity in resource availability (sequential exposure to alternative resources). Second, we quantified the induction of preference when two resources were thoroughly mixed and presented simultaneously. Third, we quantified the induction of preference under spatial resource heterogeneity, wherein alternative resources were presented to the larvae in separate patches and larvae were free to experience both resources by moving between patches. Finally, we tested the hypothesis that induction of resource preference is adaptive for larvae.

## METHODS

### Beetle stock maintenance and experimental individuals

We generated an outbred laboratory population from 12 wild-collected populations of *T. castaneum* from across India. We maintained the population at 34 °C (± 1°C) in 750g whole-wheat flour (ancestral resource; henceforth “wheat”) on a 35 day discrete-generation cycle with 2500 to 5000 individuals per generation. *T. castaneum* is a generalist and can consume a wide variety of resources. We conducted our assays using the ancestral resource for our population – whole-wheat flour (abbreviated ’W’) – and four novel resources: finger millet flour (*Eleusine coracana*, ’FM’), sorghum flour (*Sorghum bicolor*, ’S’), refined wheat flour ("*maida"* flour, ’RW’) and rice flour (*Oryza sativa*, ’R’). These are among the most common flours used in India; hence, *T. castaneum* is likely to encounter them in Indian storage warehouses. *T. castaneum* larvae perform best in their ancestral resource, wheat flour (Figure S1). They show significantly higher mortality in rice flour (∼23% drop in survival compared to wheat), and significantly delayed development in rice and finger millet flours (>100% and ∼37% delay in development rate respectively, relative to wheat) (Figure S1).

We derived all experimental individuals from generations 12-23 of the outbred stock population. To generate experimental individuals, we allowed 250-400 randomly picked stock adults (7-14 days post-eclosion) to oviposit in 150 g of finely sifted wheat flour for 24 hours. We collected eggs and placed each egg in a well of 48- well polystyrene plates (Sigma Costar; hereafter "treatment plates"), with ∼1 g of the appropriate resource per egg. We placed all plates inside a dark incubator maintained at 34 °C ± 1°C while larvae developed. Each 48- well plate contained replicate eggs for a single resource treatment. After 14 days of larval development, we tested the resource choice of each larva as described below. We tested larval choice on the 15^th^ day of development since at this stage larvae can be easily handled and observed, but still need to feed for a few days before pupating (hence their resource choice at this stage could have fitness consequences). For each experiment described below, we aimed to test an equal number of individuals per treatment. However, for various reasons (e.g. an egg did not hatch, a larva was accidentally killed during handling), the actual sample size differed slightly across treatments. For clarity, we provide the sample size for each treatment in the relevant figures in the main text and supplementary information.

### Testing larval resource choice

We measured larval resource choice in petri-plates containing adjacent patches of two resources. We spread ∼0.8 g resource on each half of a 60 mm petri-plate in a thin layer, with no space between the resources except for a circular area in the center of the plate (17 mm diameter) where we placed the 15 day old larva with no bias in orientation (Figure S2A). This placement ensured that each larva had the opportunity to sample both resources presented in the choice test. We noted the position of each larva every 12 hours (± 1 hour) for 48 hours. All larvae were housed in a dark incubator at 34 °C (± 1°C), and were outside the incubator for < 2 hours when we transferred larvae into new resources (see below) or while we noted larval position during the choice assay. For each larva, we calculated preference for the novel resource as the percent occurrence in the new resource patch. Larval tracks visible from the bottom of the petri-plate showed that larvae were capable of exploring the entire plate in a 12 hour time period. For all our experiments, these tracks confirmed that almost all larvae explore both resources, even if they showed an extreme preference for either resource. This suggests that larval movement across time points was not constrained by patch size.

Initially, we collected data on larval position for 72 hours. However, we found that some larvae from each treatment pupated towards the end of this period. Since pupae are non-feeding, data collected close to pupation may be unreliable with respect to feeding choice. For larvae that did not pupate in 72 hours, we found that larval preference calculated after 48 hours was well correlated (Spearman correlation coefficient > 0.8) with preference calculated from 72 hours of data (Figure S3). Hence, for subsequent experiments, we only collected 48 hours of data on larval position (4 data points per larva). On average, <1% of the larvae pupated during the 48 hour period across all treatments, and these were excluded from subsequent analysis.

Our assay assumes that larval presence in a resource patch indicates foraging in that patch, and that time spent foraging in a patch is proportional to the time spent feeding in that patch. If the larva was not visible during readings, we used a pair of forceps to gently move through the resource patches until the larva was found. If the larva was found in contact with both resource patches during a reading, it was noted as being present in both resources. Outside of the duration of larval transfers into new resources and the reading during the choice assay, all larvae were held in a dark incubator at 34 °C (± 1°C). Since the resources we used have different textures, we measured differences in larval mobility in each resource to account for resource-specific movement rates. We did this by measuring the number of times larvae moved across patches of the same resource in a similar setup (Figure S2B). Larval movement rate was similar across resources, suggesting that slower larval movement in novel resources does not confound our measure of larval preference for the novel resource (Figure S4).

### Analyzing data to test for induction of larval preference

To test the impact of temporal resource heterogeneity on the occurrence and magnitude of induction of preference, we exposed larvae to alternative resources either sequentially or simultaneously. To measure induction, we compared the resource preference of test larvae (given prior experience with a novel resource) to the preference of control larvae that had never experienced the novel resource (i.e. that had developed for 14 days only in wheat). A significantly higher preference for the new resource in test vs. control larvae would indicate induction of preference. For example, if rice patch occupancy of control larvae was ∼20%, but was ∼60% for larvae exposed to rice during development, this would indicate an induced preference for rice.

We conducted all statistical analyses in R (R core team, 2016) using RStudio (RStudio team, 2015). For each experimental treatment measuring larval resource choice, we had position data for four time points per individual. We measured preference for a novel resource as patch occupancy, i.e. the proportion of times (out of 4 measured) that a larva was observed in the novel resource. Since our measurements uncover strong and repeatable patterns of preference across blocks (not reported here) and we find that larvae frequently explore both patches within a 12-hour time window, this measure reflects true preference rather than sequential auto-correlation between time points. We used a generalised linear model with binomial errors and a logit link function to test whether patch occupancy differed significantly between control larvae and test larvae from the same experiment, unless mentioned otherwise. The odds ratio calculated from the fitted model coefficients is reported as the fold increase in patch occupancy in a treatment relative to the control. For example, a three-fold increase in preference (i.e. patch occupancy) for a novel resource in a treatment indicates that the larvae in this treatment were three times more likely than the control larvae to choose the novel resource.

### Testing preference induction with temporal resource heterogeneity

To test whether induction of resource preference depends on the timing of experience with a resource, we changed the resource available to larvae at different time points during development. For instance, to test the effect of early vs. late experience, we allowed larvae to experience a novel resource only during the first or the second week of development. The remaining development was completed in the ancestral resource. For this experiment, we placed eggs in a specific resource for the first week of development (as described above). To change larval exposure to a resource, we pooled week-old replicate individuals from a treatment plate and sifted them out of the resource using a fine sieve. The first resource was shaken off from their body during the sifting process. We used a fine brush to place each larva into a well of a clean 48-well plate containing the second resource. For treatments that required exposure to a single resource throughout larval development, we followed the same procedure at the 1-week time point, but placed larvae in fresh resource of the same type. We used this protocol to test the impact of larval experience with distinct resources at various time points. Therefore, we measured the impact of early (only first week of development), late (only second week of development) or continuous experience (both weeks of development) with four novel resources, on larval preference for the novel resource.

To test the impact of larval experience with finger millet immediately prior to the choice assay, we followed the above protocol, but allowed larvae to develop in the first resource for 13 days and then transferred them to the second resource for 1 day. On the following day (day 15), we measured larval preference as above. Therefore, we had four treatments – larvae that only experienced either wheat or finger millet throughout (but were transferred on day 13 to control for the impact of transfer), larvae that developed for 13 days in either wheat (or finger millet) but saw finger millet (or wheat, respectively) one day before the choice test.

### Testing specificity of the induced preference

We tested whether larval preference induced by a particular resource was specific to the experienced resource rather than a generalized acceptance of any new resource. We allowed larvae to develop in one of three treatments for 14 days: no exposure to finger millet, second week exposure to finger millet, or complete development in finger millet. On the 15^th^ day, we gave each larva a choice between the ancestral resource (wheat) and one of the four novel resources (finger millet, sorghum, refined wheat and rice). An increased preference only for finger millet would indicate that the induced preference is specific to the experienced resource and is not a general increased acceptance of any novel resource. Another possibility is that experience with a novel resource increases repulsion from the ancestral resource, which may be interpreted as increased preference for the novel resource in our assays. To test this, we allowed larvae to develop in one of four treatments for 14 days: no exposure to finger millet, only first week or second week exposure to finger millet, and complete development in finger millet. On the 15^th^ day, we gave larvae a choice between two novel resources: sorghum and finger millet. If exposure to finger millet did not increase repulsion from wheat, then larvae would still choose finger millet in this assay. These two experiments allowed us to confirm that the induction of preference is indeed a specific increased preference for the experienced resource, rather than repulsion from the ancestral wheat resource.

### Testing preference induction with homogeneous resource mixes

To measure the impact of simultaneous experience with two resources on larval preference, we weighed out equal amounts of two resources (wheat and finger millet). We mixed these together so that larvae would not be able to choose between the two resources while feeding (as in Parent et al. 2014). We collected eggs from stock beetles and allowed larvae to hatch and develop individually for 14 days in pure wheat, pure finger millet ("FM"), or the homogeneous mix (50% finger millet by weight) of the two resources. In three separate experiments, we then tested the preference of larvae from each of these treatments, given the following choice combinations - wheat versus finger millet, wheat versus the 50% mix, or the 50% mix versus finger millet. To test whether larvae were able to distinguish between small differences in finger millet concentrations in a homogeneous mix, we allowed larvae to experience the 50% homogeneous FM-wheat mix and gave them the following choices: 25% FM (in wheat, as above) versus 50% FM; 75% FM versus 50% FM; and 25% FM versus 75% FM. After experiencing a 50% mix, if larvae preferred it over other concentrations (such as 25% FM and 75% FM), this would demonstrate very high specificity of the induced preference. We confirmed that none of the resource mixes decreased larval survival, though the 75% FM mix did delay larval development (Figure S1).

### Testing preference induction with spatial heterogeneity in resource availability

To measure larval preference induction in a spatially heterogeneous patchy habitat, we directly placed eggs from stock individuals in a two-patch habitat containing the ancestral resource and a novel resource. To create the patchy habitat, we spread a thin layer of ∼1 g resource on either half of 60 mm petri-plates. We placed an egg in one patch of this habitat with a fine brush, and allowed it to develop undisturbed for 14 days (one egg per petri-plate). During development, larvae were free to move across patches and consume the resource of their choice. After 14 days, we noted the position of each larva as an overall indicator of larval preference during development and tested its resource preference as described above.

### Estimating the impact of larval choice on larval fitness

To determine the fitness consequences of experience-mediated larval resource choice, we collected eggs from the stock population and measured larval survival and development rate under conditions of temporal heterogeneity in resource availability. We exposed larvae to wheat or finger millet during the first, second and third week of development, alternating between the two resources. Larvae in control groups experienced homogeneous conditions and experienced either wheat or finger millet throughout (but were transferred to fresh flour each week). This experimental design mimicked larval resource use under spatial as well as temporal heterogeneity, if larvae acted on their resource preference. For example, while exploring a heterogeneous habitat, if a larva experienced finger millet and increasingly preferred it to wheat, it would consume largely finger millet during the remainder of its development. Larvae from the wheat-finger millet-finger millet treatment would represent this scenario, allowing us to determine the fitness impact of induced preference for finger millet relative to larvae that never experienced finger millet (i.e. the wheat-wheat-wheat controls). Note that if we measured larval fitness in a patchy habitat, we would not be able to determine which resource larvae had consumed, and would thus be unable to connect preference to fitness. After three weeks of development, we counted the number of surviving larvae and noted their developmental stage. A higher proportion of adults and/or pupae would indicate faster development, which is typically beneficial for early reproduction and reduced probability of larval parasitism (A. Fred West 1960; Blaser & Schmid-Hempel 2005). We used a generalised linear model with binomially distributed errors and a logit link function to test whether the proportion of live individuals and proportion of pupae or adults out of all live individuals were significantly different across treatments.

## RESULTS

### Induction of larval preference for new resources is time-sensitive and resource-specific

We tested whether prior experience with four new resources – finger millet, sorghum, refined wheat and rice flour – could elicit an induction of preference for these resources. We exposed larvae to one of four treatments – no exposure to the novel resource (control), exposure either during the first or second week of development, and exposure throughout development. We found that prior experience with a new resource typically increased larval preference for that resource, compared to the baseline preference shown by control larvae (Figure 1, Table S1). However, this induction of preference occurred only when larvae were exposed to the novel resource during the second week of development. In contrast, when early experience with a new resource was followed by a week in wheat flour, larvae did not show an increased preference for the novel resource. The only exception to these patterns was rice flour. Both baseline and induced preference for rice flour were very low, indicating that larvae are extremely averse to rice (Figure 1D). In contrast to the results for other resources, larvae exposed to rice flour during the first week of development showed a reduced preference for rice, although the effect size was very low (Figure 1D, Table S1).

**Figure 1:**
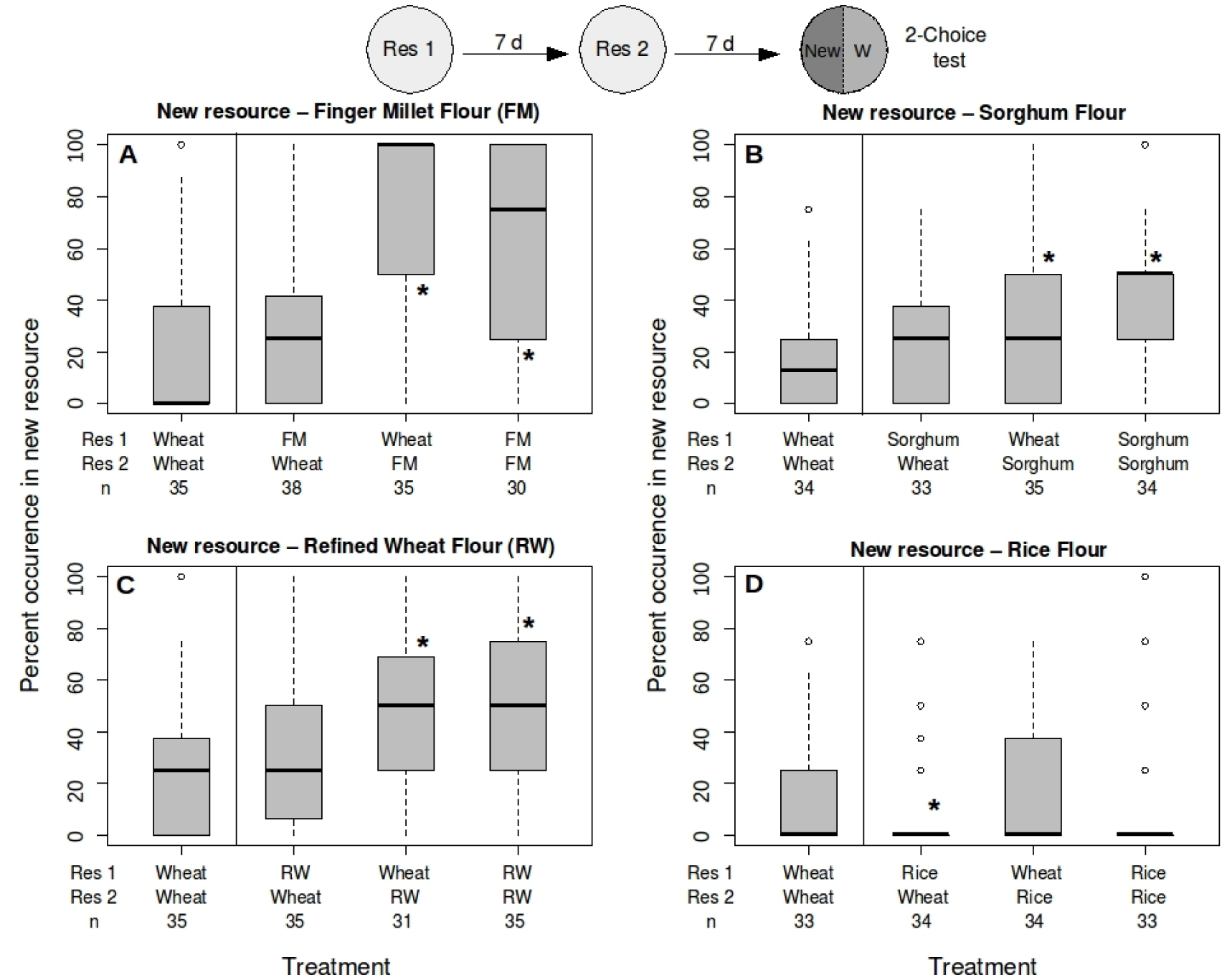
Induction of larval resource preference with two different resources presented sequentially 620 during development. Each panel shows preference of single larvae for a novel resource when given a choice 621 between the ancestral resource (wheat, "W") and the novel resource for 48 hours. Percent occurrence in a novel resource was calculated based on 4 readings of larval patch occupancy. The novel resources in each panel are (A) 623 finger millet flour (B) sorghum flour (C) refined wheat flour (D) rice flour. Larvae were reared in resource 1 for 624 the first week, and resource 2 in the second week (x-axis). In each panel, the left bar represents a control treatment where larvae were never exposed to the novel resource until the choice test. Numbers below each 626 boxplot indicate sample size. Asterisks indicate treatments that differ from the control (p < 0.05, generalized 627 linear model with binomially distributed errors). Boxplots indicate median and first and third quartiles, and 628 whiskers indicate the range of data. All figures with boxplots follow this convention.

Overall, the magnitude of the induction of preference varied across resources, with a greater than six-fold increase in preference for finger millet (Figure 1A) but a modest two-fold increase for sorghum (Figure 1B) and refined wheat (Figure 1C) (Table S1). We found that an individual’s sex did not significantly affect the magnitude of preference induction (Figure S6). Interestingly, longer exposure to the new resource (for 2 weeks) also did not alter the effect size of preference induction, except for a decrease in preference for finger millet with longer exposure (compare the last two bar plots in Figure 1A; Table S1). For the following experiments, we focused on finger millet flour, which showed the largest effect size for induction of preference.

As described above, recent (2^nd^ week only) but not early (1^st^ week only) experience with a new resource induced preference for the resource. However, it was unclear whether induction of preference was contingent on the developmental stage (i.e. the second week of larval development) or on exposure to the novel resource immediately prior to the preference test. Thus, we tested whether reducing the period of later experience could still induce preference. We found that even one day of exposure to finger millet immediately before the choice test was sufficient to induce preference, albeit with a decrease in the effect size (Figure 2, Table S2). Notably, one day of exposure to wheat was also enough to counter the effects of previous exposure to finger millet for thirteen days (Figure 2). These results strongly suggest that recent exposure to finger millet, rather than exposure during a specific developmental stage, is critical for induction of larval preference for finger millet.

**Figure 2:**
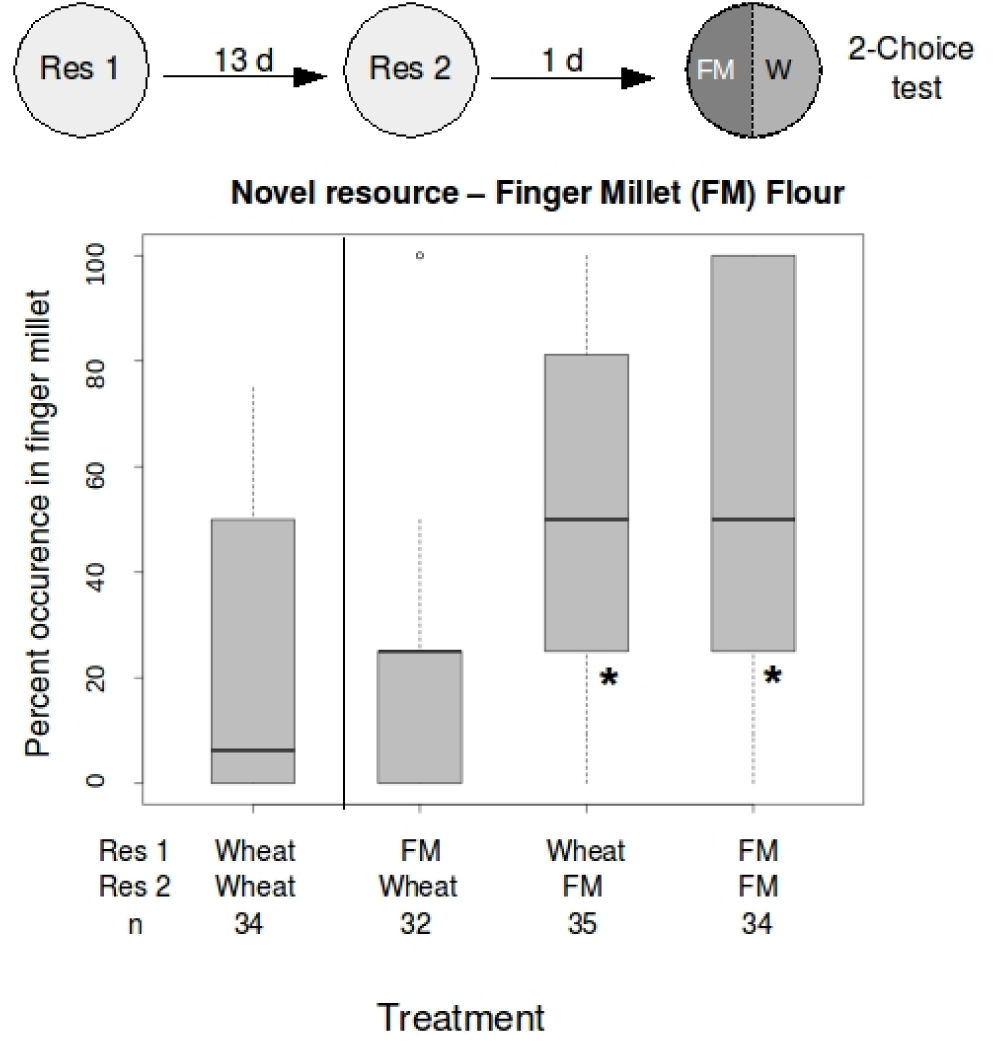
Induction of preference for finger millet occurs in a short time. Larvae were reared in resource 1 for 13 days and then resource 2 for 1 day (x-axis). Larval preference was measured as described in Fig 1.Boxplots represent data as described in Figure 1.

Next, we tested whether the induction of larval preference for finger millet flour is specific to finger millet, or is generalized to other new resources. We found that induction of preference was generally specific to the experienced resource (Figure S7, Table S3), although we did observe a small increase (∼1.8 fold) in preference for refined wheat flour in larvae exposed to finger millet in the second week of development. Finally, we tested whether increased occupancy of the finger millet patch in the choice test (Figure 1A) represents repulsion from wheat flour (ancestral resource) or attraction to finger millet. After the larva had experienced finger millet, we offered it a choice between finger millet and sorghum. We found that the induction of preference for finger millet was intact in this novel context, demonstrating that aversion to wheat flour does not explain the induction of preference observed in finger-millet flour. Taken together, our results demonstrate a specific, experience-mediated increase in preference for novel resources in flour beetle larvae (Figure S8, Table S4).

### Simultaneous experience with two resources also elicits specific induction of larval preference

In the above experiments, we presented larvae with alternative resources in a temporally distinct manner (i.e. only one resource at a time). Does induction of preference occur when larvae experience multiple resources simultaneously, and how specific is the induced larval preference under such conditions? To test this, we exposed larvae to a homogenous mixture of finger millet flour and wheat flour (1:1 by weight) for fourteen days, with exposure to pure wheat and pure finger millet flour as controls. On the fifteenth day, we tested whether larvae from each regime preferred 100% wheat, 100% finger-millet flour, or a 1:1 homogenous mixture of both flours (henceforth ’mix’). We found that larvae strongly preferred the same resource (or mix) that they had experienced previously (Figure 3, Table S5). Notably, larvae that were first exposed to the 1:1 mix and then presented with the two pure resources in the choice test showed a strong preference for pure finger millet flour (Figure 3A). These results suggest that the presence of the ancestral resource does not qualitatively interfere with preference induction for the novel resource, although the effect size was about half that observed for induction with pure finger millet (Figure 3A; Table S5A). Interestingly, larvae reared on a 1:1 finger millet-wheat mix did not distinguish between mixes with different proportions of finger millet and wheat. For example, when given a choice between the 1:1 mix versus a 25% or a 75% mix, larvae that developed in the 1:1 mix did not show a preference for the experienced mix (Figure S9). Together, these results show that even half the amount of finger millet experienced simultaneously with wheat can induce preference for finger millet; but larvae are unable to distinguish quantitatively between resources mixes with different proportions of finger millet.

**Figure 3:**
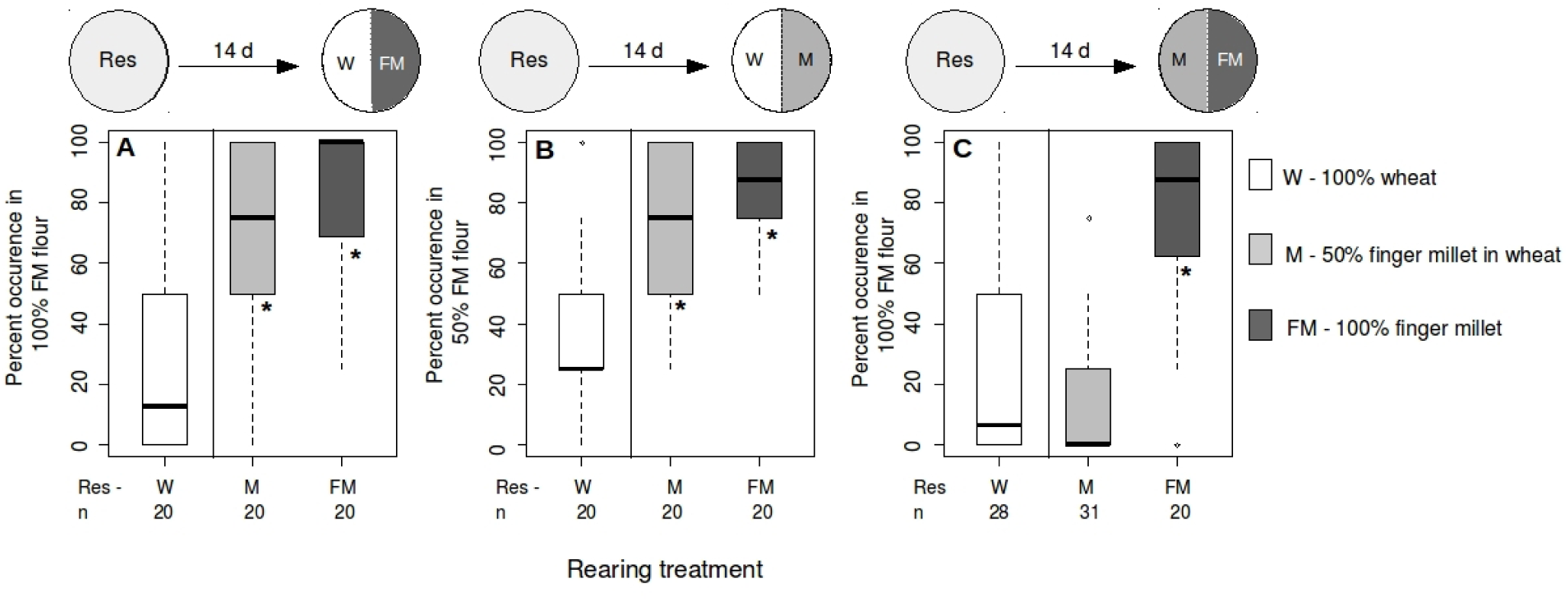
Induction of preference also occurs with a mixture of finger millet and wheat flour. Each panel shows preference of single larvae for a resource mix with the higher concentration (by mass) of finger millet flour when given a choice between various combinations among 100% wheat flour (white), 1:1 mix of finger millet flour and wheat (light grey) and 100% finger millet flour (dark grey) for 48 hours. Larval preference was measured as described in Figure 1. The choice combinations in each panel are (A) 100% wheat versus 100% finger millet (B) 100% wheat versus 50% finger millet and (C) 50% finger millet versus 100% finger millet. Larvae were reared in each treatment (x-axis) for 14 days. Boxplots represent data as described in Figure 1.

### Impact of spatial habitat heterogeneity on induction of preference

Next, we tested whether a spatially heterogeneous habitat can alter larval preference via induction. We placed individual eggs in one patch of a two-patch habitat containing the ancestral wheat resource, and either finger-millet ("FM") flour or refined wheat ("RW") as a new resource. Larvae developed while moving freely across patches, consuming their resource of choice. After 15 days, we found that in the FM-wheat habitat, larvae typically occupied the same resource patch where they were placed as eggs (i.e. the natal resource; Figure 4A). Regardless of where they were placed as eggs, larvae that developed in the patchy FM-wheat habitat showed a consistently higher probability of choosing finger millet in the choice test (∼40%; Figure 4B). Thus, prior contact with the finger millet even during exploration of the habitat was sufficient to induce a preference for finger millet. In contrast, in the RW-wheat habitat, most individuals that were initially placed in the refined wheat patch moved to the wheat patch over the course of development (Figure 4A). However, larvae initially placed in refined wheat did show a significantly higher preference for refined wheat compared to individuals placed in the wheat patch as eggs (Figure 4C). In this habitat, it appears that the natal resource induced larval preference. Hence, the relative use of the natal resource in a patchy habitat varies depending on the alternative resource available in that habitat.

**Figure 4:**
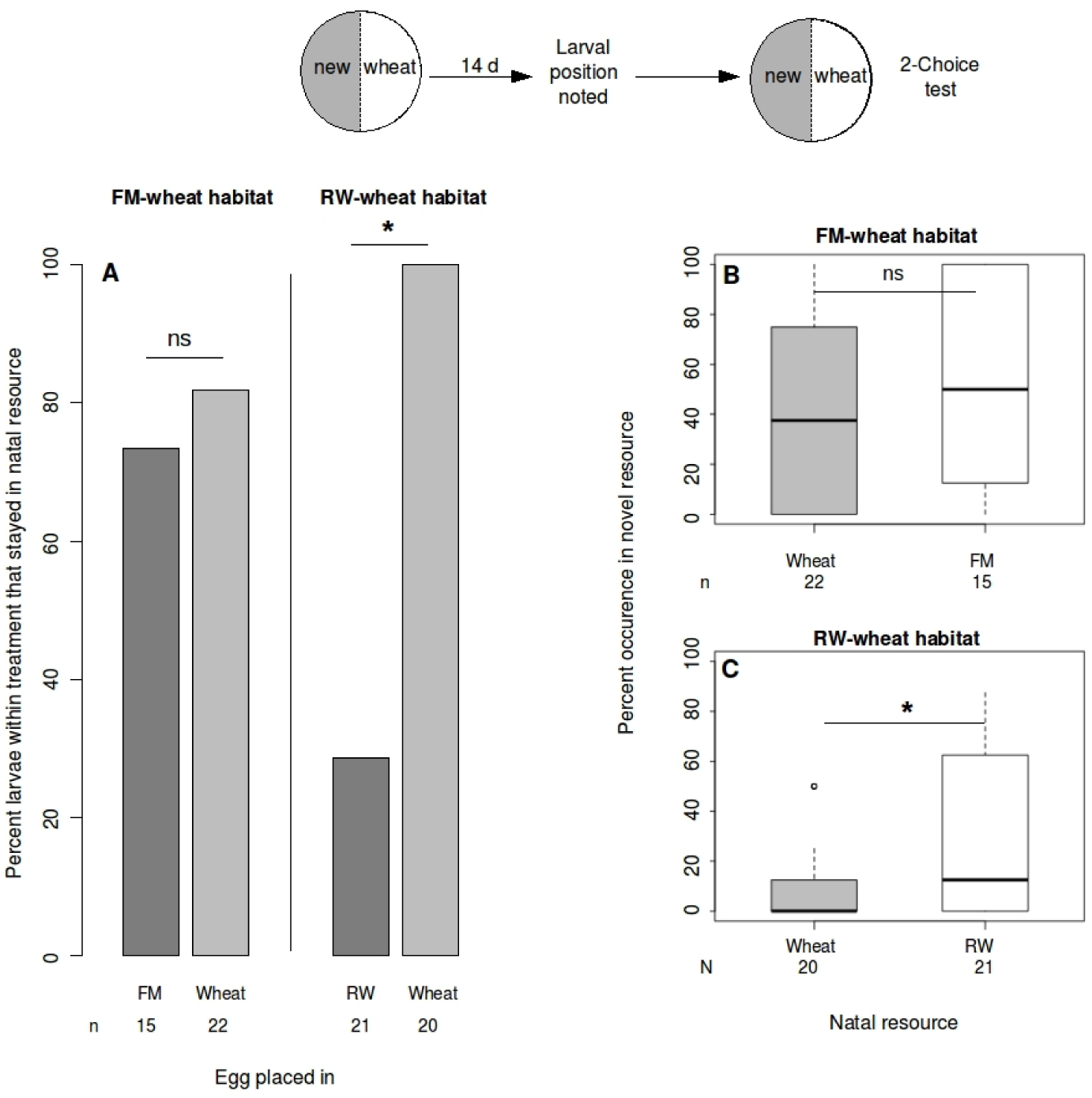
Larval preference for new resources after development in a heterogeneous two-patch habitat. Individual eggs were placed in one patch of a two-patch heterogeneous habitat, and larvae were allowed to develop allowing for unrestricted movement across patches. (A) Percentage of larvae found in the same patch they hatched into (natal resource) in either an FM-wheat (left) or an RW-wheat (right) patchy habitat after 14 days of development. Asterisks indicate a significant difference between the indicated treatments (p < 0.05, pairwise two-sided chi-squared test of proportions with Yates’ continuity correction). The new resource in subsequent preference tests on the 15^th^ day was either (B) finger millet (FM) or (C) refined wheat (RW) flour. Larvae within each treatment are split along the x-axis according to their natal resource. The y-axis denotes larval preference for the natal resource. Asterisks indicate a significant difference in percent occurrence in the new resource across individuals in different natal resources. (p < 0.05, generalised linear model with binomially distributed errors). Boxplots represent data as described in Figure 1.

### Induction of preference can be maladaptive for larval development

What are the fitness consequences of larval preference induction? Earlier, we found that larvae that developed in a new resource did not pay a mortality cost, except in rice where larval mortality was significantly higher than in wheat (∼65% survival in rice, ∼80% in wheat; Figure S1). Finger millet and rice were both suboptimal for larval development rate (Figure S1). Of the four tested resources, the only resource that did not induce preference was rice flour, which is also the only resource that substantially decreases larval survival. However, we found that across resources, the magnitude of preference induction was not associated with larval development rate in those resources.

In the experiments described above, we measured the fitness consequences of consuming a single resource throughout larval development. To simulate the fitness consequences of varying larval resource choice in a heterogeneous habitat, we transferred the larvae to a different resource for three developmental windows of one week each. This represents a scenario in which larvae move across resource patches every week. Thus, we could estimate the fitness consequences of larvae exerting preference and consuming different preferred resources over the course of their development. For example, to estimate the development rate of larvae that experienced finger millet in the second week of development in a heterogeneous habitat (containing wheat and finger millet patches), we allowed larvae to develop in wheat for one week, finger millet for the second, and wheat for the third. After allowing larvae to develop in these regimes, we measured the fraction of larvae that successfully pupated or eclosed as adults as a fitness proxy.

We found that regardless of the treatment, nearly all larvae pupated within three weeks, and had similar rates of survival (Figure S10). Of the larvae that developed in wheat for the first two weeks of development, ∼10% had also completed the pupal stage and eclosed as adults (Figure 5). However, almost none of the larvae that experienced finger millet during the first two weeks had eclosed as adults, showing significantly delayed development. These data suggest that an early preference for finger millet could delay larval development, suggesting that induction of preference for finger millet may be maladaptive for larvae.

**Figure 5:**
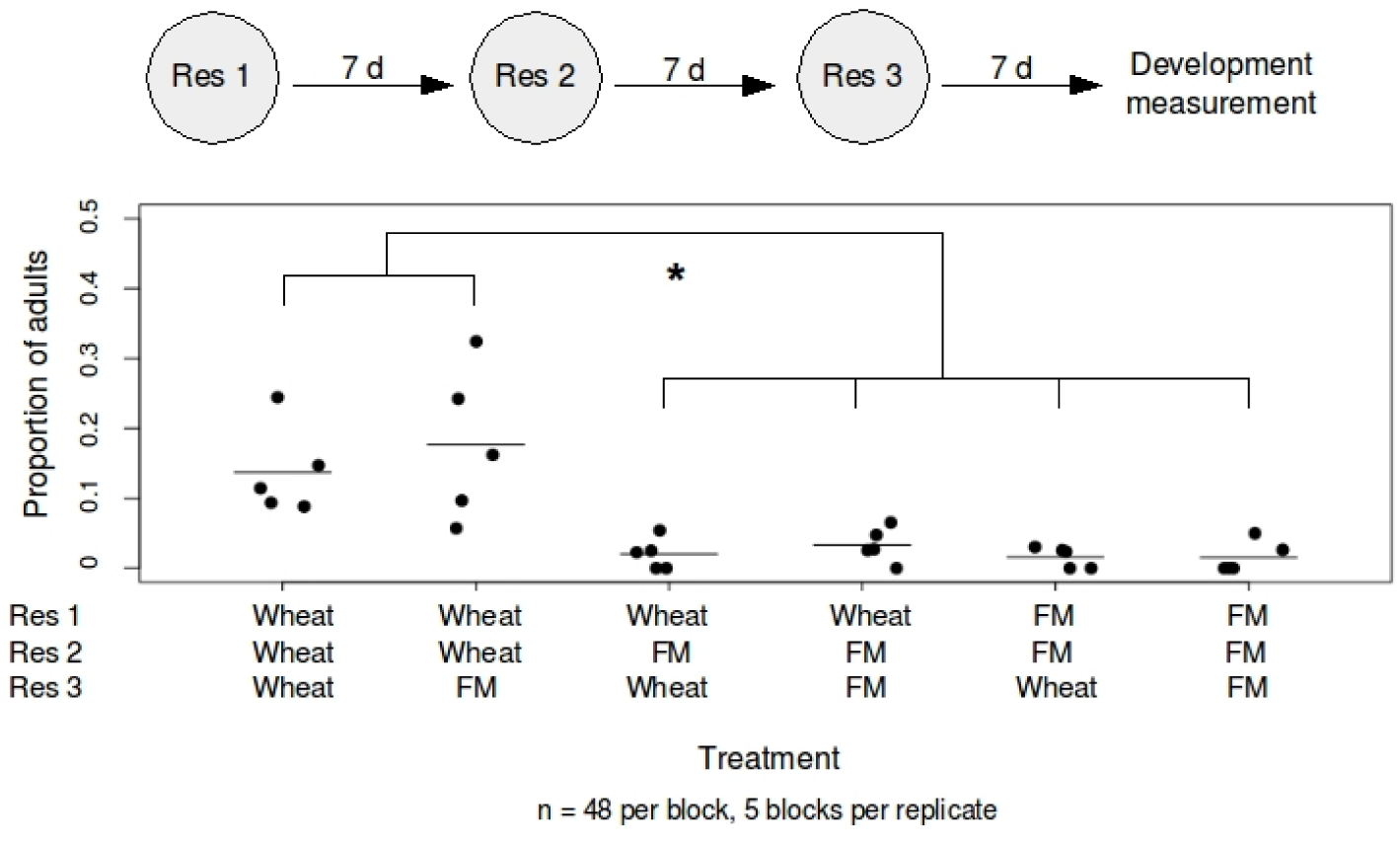
Development rate of larvae as a function of their respective developmental experience. Each 659 point represents data for a single experimental block. Within each block, individual eggs (n = 48 per 660 developmental regime) were placed in each of the treatments represented along the x-axis. The y-axis represents 661 the proportion of individuals that had eclosed as adults on the 21^st^ day. Lines within the scatter represent mean of 662 the data. Asterisks indicate significant difference in development rate between treatments as represented (p < 663 0.05, generalized linear model with binomially distributed errors)

## DISCUSSION

Despite a large body of work on the induction of larval preference, we understand very little about the ecological significance of preference induction. One reason is the perception that larval preference is ecologically irrelevant, and that adult oviposition preference largely dictates larval performance in most herbivorous insects (Dethier 1959; Jaenike 1978). However, many studies have also emphasized larval participation in host choice – especially in highly heterogeneous habitats with multiple resources (Wiklund 1975; Chew 1977; Berdegue et al. 1998; Mayhew 2001; Gamberale-Stille et al. 2014; Abbott & Dukas 2016) – and in determining their own fitness at finer spatial scales (Soler et al. 2012; Gamberale-Stille et al. 2014; Abbott & Dukas 2016). Here, we analyzed the impact of experience with resource heterogeneity on larval preference for new resources in *Tribolium castaneum*, a generalist pest. Our data show that red flour beetle larvae can exert a strong, but flexible, resource choice. We demonstrate that *T. castaneum* larvae fed on their ancestral wheat resource prefer wheat, but are more likely to choose a novel resource if they experienced it recently. The magnitude of this induction of preference for new resources varied across the four new resources that we tested. Preference induction is very specific and occurs even when alternate resources are presented simultaneously. Interestingly, induction of preference occurs readily for resources that delay larval development, but not for the one resource that substantially decreases larval survival. Thus, we show that induction of preference can play a powerful role in determining larval resource choice and altering their fitness in complex environments.

Our finding of resource-specific effect size of preference induction corroborates results from earlier studies with Lepidopteran larvae (Jermy et al. 1968; de Boer & Hanson 1984; Boer 1992; Silva et al. 2014), and suggests that such resource specificity in preference induction may be a general phenomenon. The variable magnitude of induction may be related to the mechanism underlying preference induction, which is partially characterized in only a few insects. In Lepidopterans such as *Manduca sexta* and *Pieris rapae,* as well as locusts, dietary experience can impact the peripheral sensory system by changing the sensitivity of gustatory receptor neurons (Abisgold & Simpson 1988; Zhou et al. 2009; Glendinning et al. 2015). It is also possible that the abundance of specific compound(s) used by larvae for food recognition (del Campo et al. 2001) varies across resources, altering short-term recognition of these resources. The induction of preference with recent, brief experience (1 day), and the observed resource specificity are both consistent with this hypothesis. Therefore, we speculate that the chemical composition of the different resources may determine the strength of preference induction.

An important feature of our experimental system was our ability to test the impact of experience with simultaneously experienced alternative resources. In phytophagous insects such as Lepidopterans that directly feed on plants, it is difficult to present a mix of different resources at the same time. *A priori*, we expected that experiencing a novel resource as part of a resource mix should weaken the induction of preference, compared to experiencing a pure novel resource. Our findings are consistent with this expectation. We observe that larvae discriminate not only between pure resources but also between a homogenous mix of these resources, preferring exactly what they experienced when it is available. This suggests that when two resources are experienced simultaneously in a mix, a strong preference is induced for a mix of these resources, over a pure form of either. Interestingly, larvae that experienced a balanced mix of a novel resource (finger millet) and the ancestral resource (wheat) strongly preferred 100% finger millet over 100% wheat. A simple expectation is that larvae should not discriminate between wheat and finger millet, having experienced an equal amount of both during development. In contrast, our results suggest that different components in a single mix have quantitatively different effects on larval preference, with the ancestral resource in the mix impacting larval preference much less than an equal mass of the novel resource. Thus, some novel resources (in this case, finger millet) can induce larval preference more strongly than the ancestral resource, indicating an asymmetry in the impact of different resources on insect preference. A similar phenomenon was observed in *Manduca sexta* larvae, where across choice combinations, acceptable non-host plants induced a stronger preference than host plants (Boer 1992). However, in this study, alternative hosts were presented separately and preference was not measured using choice assays. Interestingly, we found that the specificity of induced resource preference did not extend to the exact concentrations of the experienced mix (Figure S9). Our data do not distinguish between the alternate possibilities of whether larvae are incapable of distinguishing between different concentrations of resources, or whether they do not discriminate between them. While experience with homogeneous mixes is probably not very common in nature, our data hint at the underlying mechanism of induction of preference in *T. castaneum* larvae, and are consistent with the hypothesis of additive resource-specific modifications in the sensory system leading to increased preference for a resource mix. However, we need further investigation of the exact mechanism of the induction of preference (Bernays & Weiss 1996; Anderson & Anton 2014).

An intriguing and novel finding from our work is that resource-specific induction of preference can also occur in a spatially heterogeneous habitat. Importantly, in this experiment, we found that larvae are capable of moving out of their natal resource, thereby impacting their own fitness. An earlier study with fruit flies also showed that larvae could move away from a suboptimal natal habitat chosen by their mother (Abbott & Dukas 2016). However, in our experiments, larvae did not move away from finger millet, even though it significantly delays larval development. On the other hand, larvae avoided refined wheat, which confers similar larval survival and development rate as wheat. Similarly, previous work showed that flour beetle larvae reared in a wheat-corn patchy habitat preferred the suboptimal corn resource more strongly than larvae reared in a homogeneous wheat habitat (Parent et al. 2014). Our current results suggest that this suboptimal preference for corn may have been induced by prior exposure to corn. Together, these observations indicate that alternative resources experienced at a small spatial scale may alter larval preference, and possibly the fitness of developing larvae. Thus, our experiments provide new insights about the sensitivity of larval preference to their natal environment and nearby alternative resources and underscore the importance of studying larval preference in heterogeneous habitats.

By explicitly measuring larval resource preference, movement and fitness, we showed that larval resource use is strongly driven by resource-specific induction rather than the potential fitness consequences of their choice. On the one hand, the lack of induction with rice flour may be seen as adaptive, since larval survival is significantly lower in rice compared to the other resources (all of which showed significant preference induction; Figure S1). On the other hand, larval development rate was not correlated with the strength of preference induction across resources. For example, larval preference was most strongly induced by finger millet, a resource that decreases development rate substantially. These results are similar to previous findings in *Manduca sexta,* where the strength of larval preference induction was not correlated with larval performance across hosts (Silva et al. 2014). If beetle larvae acted on induced preference during early development, they would suffer a significant developmental delay (∼10-18% lower eclosion rate at three weeks) compared to larvae that did not consume finger millet. Given these fitness consequences, optimally behaving larvae should move out of a finger millet patch, particularly during the first two weeks of development. However, our data show that this does not occur in a patchy habitat, possibly due to the strong induction by finger millet. Taken together, our experiments suggest that the induction of resource preference may be maladaptive, and may serve as a barrier to optimal resource choice in heterogeneous habitats.

Is the induction of larval resource preference broadly relevant in insects? In the specific case of flour beetles, habitat heterogeneity and resource turnover is likely a common feature in stored grain warehouses, particularly where the parent populations used in this study were collected. However, we could not systematically measure temporal or spatial heterogeneity in the natural habitat of *Tribolium castaneum*, and we hope that future studies can quantify such heterogeneity. More generally, if insect larvae explore their habitats, they may often encounter local heterogeneity in patch quality (as in Abbott & Dukas 2016; Soler et al. 2012; Gamberalle-Stille et al. 2014). Our data suggest that larval resource use in such heterogeneous habitats can be driven by several features of the habitat, such as spatial and temporal heterogeneity in resource availability. In this context, our experiments provide new insights about the sensitivity of larval preference to their environment and underscore the importance of studying larval preference in heterogeneous environments. Larval resource choice may also play an important role in the context of suboptimal maternal oviposition. A growing body of evidence suggests several constraints on optimal female oviposition in generalists (Charlery de la Masselière et al. 2017), such as search time, patch complexity, and the ability to receive and process sensory information (Janz & Nylin 1997; Mayhew 1997; Bernays 1999; Bernays & Funk 1999; Bernays 2001; Janz 2002). Thus, studying only oviposition preference in such habitats may not reliably predict larval resource use, and alternative resources experienced at a smaller spatial scale may alter the fitness outcomes for developing larvae.

In closing, we note that many questions regarding the ecological and evolutionary consequences of larval resource choice remain unanswered. This is primarily because larval choice has been underrepresented in studies of insect host choice. In general, studies of host preference have largely focused on Lepidopterans, which have a unique life history compared to other insect orders. More studies of insect host use across insect life stages and in non-Lepidopteran species may help us develop general principles of insect host choice and understand its implications for individual fitness. Experience based plasticity in larval resource choice may be an important mechanism through which rapid behavioural diet shifts can occur, either when female oviposition behaviour is suboptimal, or under conditions of habitat heterogeneity where larval resource availability fluctuates. Under such conditions, the induction of preference may enhance larval performance on suboptimal resources through increased feeding (Agrawal et al. 2002b). We find that induction of preference does not occur for a resource that substantially decreases larval survival. Therefore, this may well be a mechanism through which rapid niche expansion can occur within the larval stage of individuals, altering subsequent acceptability of the novel resource and allowing individuals to utilize and adapt to novel resources (West-Eberhard 1989). Preference induction may thus facilitate local adaptation in populations experiencing heterogeneity in resource availability. Our study represents the first systematic analysis of the impact of habitat heterogeneity on larval preference for new resources, and suggests its potential role in the evolution of diet breadth.

## Acknowledgements

We thank members of the Agashe lab and Shannon Olsson for discussion and comments on the manuscript. This work was supported by the National Centre for Biological Sciences and an INSPIRE Faculty award to DA (IFA- 496 13 LSBM-64).

## Author contributions

DA, VRK and SI designed experiments; VRK and SI conducted experiments; VRK, SI and DA analyzed data;

VRK and DA wrote the manuscript. All authors gave final approval for publication.

